# mScarlet3-H with low brightness and fluorescence lifetime has potential for cellular lifetime-unmixing and lifetime-based pH-sensing applications

**DOI:** 10.1101/2025.07.15.664898

**Authors:** Theodorus W.J. Gadella, Laura van Weeren

## Abstract

An independent evaluation of the spectroscopic properties, cellular performance, merits and pitfalls of the red fluorescent protein mScarlet3-H as compared to mScarlet3 is reported. mScarlet3-H was generated from mScarlet3 by a single M163H mutation. Purified mScarlet3-H is characterized by a molar coeWicient of 79,040 M^-1^cm^-1^, afluorescence quantum yield of 17.8%, molecular brightness of 14.1 and a heterogeneous multiexponential decay with an average fluorescence lifetime of 1 ns. Evaluation in living mammalian cells revealed a comparable maturation speed and eWiciency of mScarlet3 and mScarlet3-H, but the overall cellular brightness of mScarlet3-H was 5-fold lower than that of mScarlet3. Photobleaching analysis in live cells revealed identical photobleaching kinetics of mScarlet3-H and mScarlet-H. Thefluorescence intensity,fluorescence spectra and fluorescence lifetime of mScarlet3-H were found to be strongly pH-dependent between pH 4-8. The fluorescence lifetime increased from 1 ns to 3 ns in lowering the pH from 8 to 4 with a pK of ∼6. The much lower lifetime of mScarlet3-H (∼1 ns) as compared to mScarlet3 (∼4 ns) allows dual fluorescence lifetime unmixing applications in single channel FLIM recordings in compartments with neutral to slightly alkaline pH. Furthermore, the strongly pH-dependent fluorescence lifetime of mScarlet3-H enables fluorescence lifetime-based pH sensing in a pH region between pH 5 to 7. With this property autophagy of the cytoplasm can be visualized by the pH-dependent fluorescence lifetime with mScarlet3-H accumulation in lysosomes. Potential useful applications and pitfalls regarding the special properties of mScarlet3-H are discussed.

## Introduction

Redfluorescent proteins have emerged as indispensable tools for cell biology. With a 70-150 nanometer redshift as compared to popular green fluorescent proteins they allow live cell imaging at much milder conditions (Waldchen et al., 2015). The green to yellow excitation light is much better tolerated by cells and produces vastly reduced autofluorescence levels as compared to blue excitation required for exciting green fluorescent proteins. The first RFP published was DsRed (Matz et al., 1999) which was quite bright but unfortunately also an obligate tetramer. Much eNort has been devoted since to monomerizing and optimizing RFPs. Breaking up the tetramer greatly reduced the quantum yield and maturation efficiency of the ensuing monomerized RFP’s (Campbell et al., 2002). A popular mRFP was mCherry that displayed decent fluorescence intensity, maturation and photostability, however it had a quantum yield of only 20% (Shaner et al., 2004). In 2016 we published the generation of a much brighter monomeric RFP mScarlet (Bindels et al., 2017) evolved from a completely synthetic DNA template. Together with mScarlet also two other variants were described: mScarlet-I, carrying a single mutation T74I improving maturation speed and -extent, but with reduced quantum yield; and mScarlet-H carrying a single mutation M164H vastly improving photostability but also vastly reducing quantum yield and brightness (Bindels et al., 2017). In 2023 after several years of further optimization we published mScarlet3, which currently is the brightest monomeric red fluorescent protein with complete and fast maturation. mScarlet3 has a quantum yield of 75%, an extinction coefficient of 104,000M^-1^cm^-1^ and it displays a mono exponential decay with afluorescence lifetime of ∼ 4 nanoseconds (Gadella et al., 2023). Recently, a study was published describing mScarlet3-H which also carries the single amino acid change M163H (Xiong et al., 2025). As a result of this mutation a marked increase in photostability was described, mimicking the earlier described effect of the same mutation in mScarlet. In an accompanying news and views paper Campbell stated that we ‘*did not incorporate this mutation into the brighter mScarlet 3*’ (Campbell, 2025). Yet this statement is incorrect because this variant was generated already in 2022 by us and it was characterized in detail, but we regarded it as a downgrade in view of its much-reduced brightness, pH instability and no increase in photostability as compared to mScarlet-H. For that reason, we decided to not include it in the paper describing mScarlet3 with applications for live cell imaging, FRET and protein tagging in order not to confuse the scientific community with partially compromised fluorescent proteins. In this paper we describe our own measurements and comparison of mScarlet3 to mScarlet3-H, as well as possible uses of its reduced fluorescence lifetime and pH instability in fluorescence lifetime unmixing and quantitative lifetime-based pH sensing in acidic cellular organelles.

## Results and discussion

### Spectroscopic analysis

Following recombinant protein production and purification, we have thoroughly measured the spectral properties of mScarlet3-H as compared to mScarlet3. Like mScarlet3, mScarlet3-H is a genuine red fluorescent protein yet with blue shifted excitation properties as compared to mScarlet: see figure 1a and 1b for absorbance, excitation and emission spectra. The spectra we measured are similar to what was published by (Xiong et al., 2025). We observe an excitation maximum at 551 nanometer (18 nanometers blue shifted as compared to mScarlet3) and an emission maximum at 582 nanometers (10 nanometer blue shifted as compared to mScarlet3). We determined the extinction coefficient using alkaline denaturation (figure 1c,d). These data indicate an extinction coefficient of 104,000 M^-1^cm^-1^ for mScarlet3 and 79,040 M^-1^cm^-1^ for mScarlet3-H. From the raw absorbance spectra in figure 1c,d, it is clear that the molecular extinction of the red fluorescing form in PBS buffer is lower for mScarlet3-H than for mScarlet3 (to yield an equal amount of alkaline denatured form with an extinction coefficient of 44,000 M^-1^cm^-1^) (Gross et al., 2000; Shagin et al., 2004). The extinction coefficient of 79,040 M^-1^cm^-1^ for mScarlet3-H is lower as compared to the value reported by (Xiong et al., 2025) but is in line with reduced extinction coefficients of other mScarlet variants carrying the M164H mutation such as mScarlet-H (64,000 M^-1^cm^-1^) (Bindels et al., 2017) and mScarlet-H-S220A (69,000 M^-1^cm^-1^) (Gadella and van Weeren, unpublished). We measured a quantum yield for mScarlet3-H of 17.8% using mScarlet3 as reference. This value compares favorably with the value reported by (Xiong et al., 2025). The both lower extinction coefficient and quantum yield result in a much-reduced intrinsic brightness value of 14.1 for mScarlet3-H, being 5.5 times lower as compared to 78.1 of mScarlet3 (see Table I).

**Figure 1.**
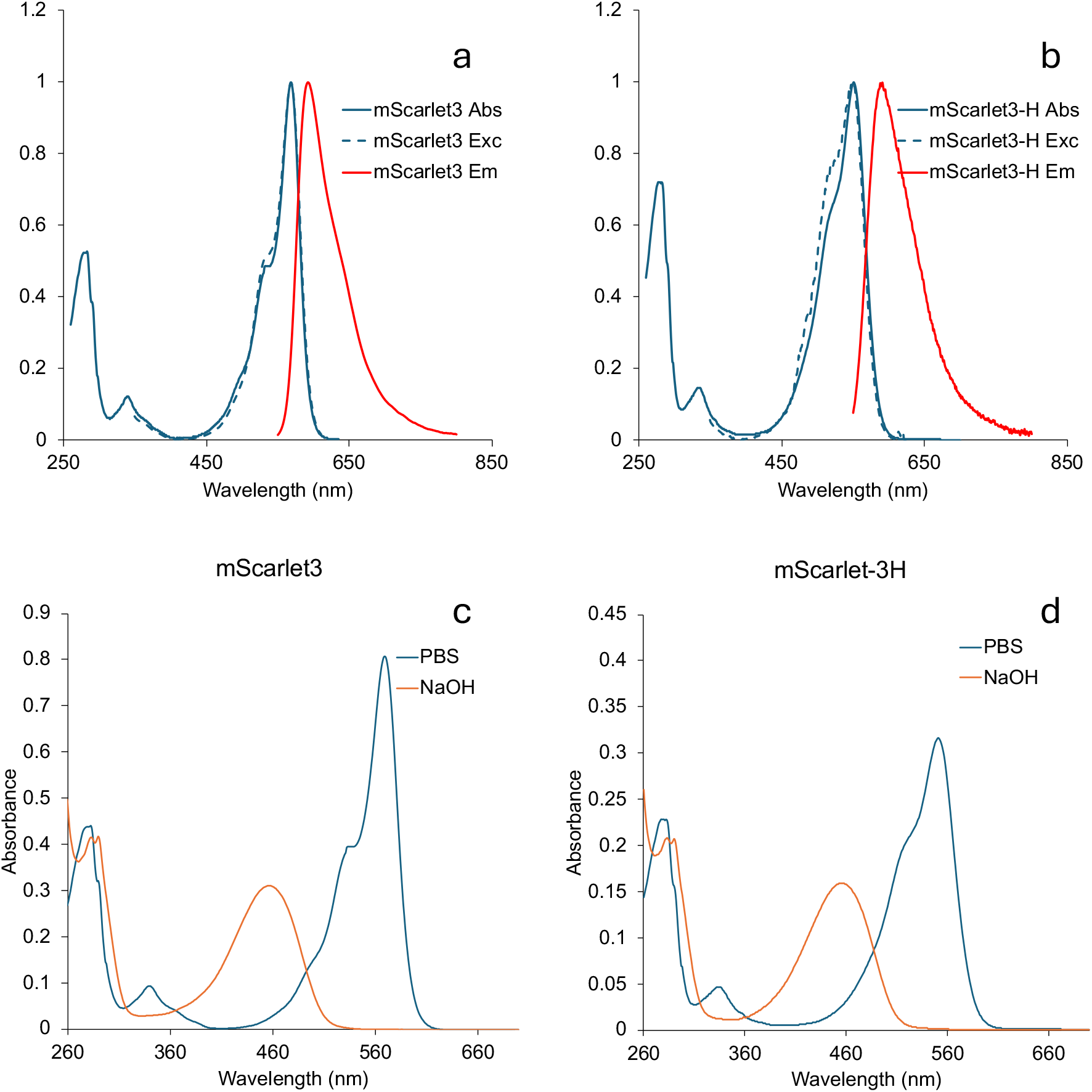
Comparison of spectral properties of mScarlet3 and mScarlet3-H. **a, b**. Normalized absorbance, excitation, and emission spectra of mScarlet3 (**a**) and mScarlet3-H (**b**). For the fluorescence excitation spectra, the emission was set at 620 nm, for the fluorescence emission spectrum excitation was set at 540 nm. **c,d.** change in absorbance spectra of mScarlet3 (**c**) and mScarlet3-H (**d**) upon alkaline denaturation.

**Table I.** Properties of mScarlet3 mScarlet3-H*.

| | Abs max (nm) | Em max (nm) | $\epsilon_{max}$ M<sup>-1</sup>cm<sup>-1</sup> | QY (%) | $\tau$ (ns) | Mol. Brightness | Cell. Brightness (%) | Maturation delay (h) | Maturation extent (%) | pK |
| --- | --- | --- | --- | --- | --- | --- | --- | --- | --- | --- |
| mScarlet3 | 569 | 592 | 104000 | 75.1 | 3.96 | 78.1 | 100 | 0.6 | 95 | 4.5 |
| mScarlet3-H | 551 | 582 | 79040 | 17.8 | 1.0 | 14.1 | 20 | 0.5 | 104 | 4 and 6 |

\*Values for mScarlet 3 are from (Gadella et al., 2023), values of mScarlet3-H are from this study.

### Brightness and maturation in cells

The cellular brightness in mammalian cells of mScarlet3 and diverse other RFPs was estimated by transfecting with a polycistronic vector, which leads to a 1:1 stoichiometric coproduction of mTurquoise2 as reference and an RFP (Bindels et al., 2020). Here the measured cellular intensity obtained in the red channel can be compared with the cyan channel for each individual cell to correct for expression level differences. By plotting the red versus the cyan fluorescence average intensity for each cell, the slope of the trendline fitting best to all data points is an accurate estimate of the raw cellular brightness (see figure 2a and figure 2b). This raw value must be corrected for the relative spectral throughput of the microscope for different RFPs. This is necessary because certain RFP’s have shifted excitation spectra yielding more efficient excitation at the employed excitation band where other RFP’s have shifted emission spectra that better overlap with the employed emission bandpass. The employed correction factor is the relative excitation efficiency (percent of maximum) and the throughput of the emission filter (percent of total emission spectrum). In our instrument we have an effective excitation bandpass between 543-558 nm and an emission bound pass between 570-616 nanometer. This yields a correction factor relative to mScarlet of 0.985 for mScarlet3 (Gadella et al., 2023) and 0.692 for mScarlet-3H. By multiplying the RFP channel intensities of mScarlet3-H, with 0.692/0.965, the normalized cellular brightness can be estimated (see Figure 2c). Hence, at employed microscopy settings, mScarlet3-H is relatively more efficiently detected than mScarlet3 (a factor 1.42 times, mainly caused by its blue-shifted excitation). After correction for spectral through-put, this yields a cellular brightness of mScarlet3-H of 20% as compared to mScarlet3, i.e. a 5-fold reduced brightness in cells 24 hours after transfection (Figure 3a). Also included is a brightness comparison by using 570 nm excitation within the same setup in another experiment. This is favoring mScarlet3 over mScarlet3-H in excitation in view of the red-shifted excitation spectrum of mScarlet3. As can be seen in comparing Figure 2d and 2e, such conditions yield a ∼12-fold higher mScarlet3 (raw estimated cellular) brightness as compared to mScarlet3-H. This high difference is expected and underlines the importance of proper spectral correction to estimate the cellular brightness and maturation extent in cells. By correcting for the poor excitation at 570 nm of mScarlet3-H relative to mScarlet3, a similar corrected relative cellular brightness can be computed as for the 550 nm excitation conditions (compare figure 2c with figure 2f). The cellular brightness of mScarlet3-H of only 20% as compared to mScarlet3, is 70% lower than the value of ∼36% reported by (Xiong et al., 2025). Since the imaging conditions with which this was measured were not indicated by Xiong et al., (Xiong et al., 2025), we hypothesize that their values might represent uncorrected raw ratiometric values obtained at (possibly blue-shifted) imaging conditions that are relatively advantageous for mScarlet3-H as compared to mScarlet3. Similarly, tdTomato was estimated to be brighter than mScarlet3 by (Xiong et al., 2025), in contrast to our earlier published findings for dTomato which is less bright than mScarlet3 (Gadella et al., 2023). In view of the blue-shifted absorbance spectrum of (t)dTomato (Shaner et al., 2004), that other discrepancy, may represent a similar bias and cause. As can be seen in Figure 2, uncorrected cellular brightness measurements can vary significantly for different imaging conditions.

**Figure 2:**
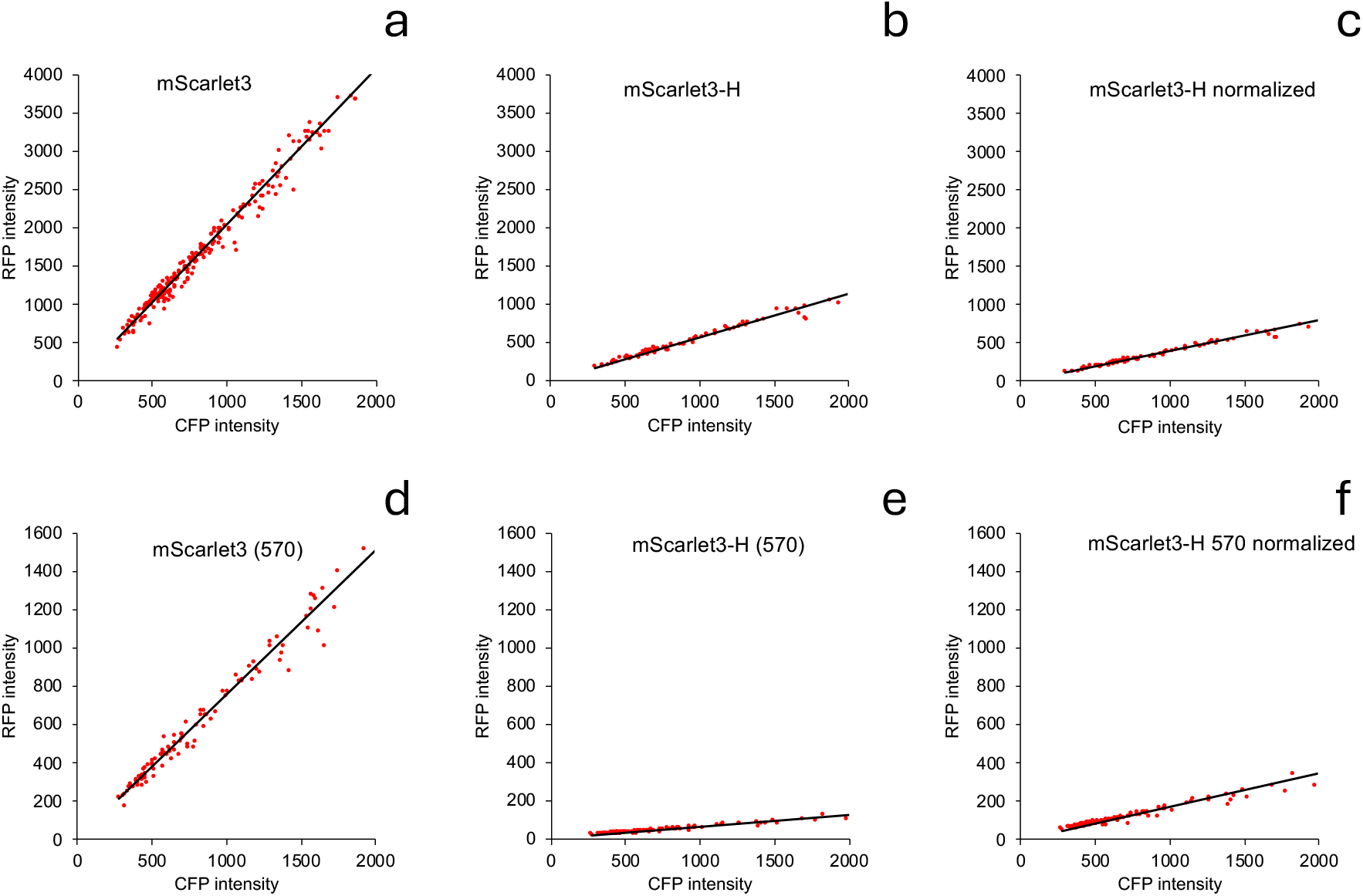
Brightness of mScarlet3 and mScarlet3-H in HeLa cells 24 h after transfection. Both mScarlet3 and mScarlet3-H were coexpressed with mTurquoise2 in HeLa cells at a 1:1 molecular ratio. Redfluorescence was detected at 543-558 nm excitation and 570-616 nm emission; cyanfluorescence was detected at 430-450 nm excitation and 459-490 nm emission**. a, b.** The average redfluorescence and cyanfluorescence was quantified for each cell (red data points, n=251 for mScarlet3 (**a**) and n=98 for mScarlet3-H (**b**)). **c** The same data for (b) are replotted after correction for spectral throughput of the microscope (correcting for instrument-dependent excitation and detection ebiciencies, relative to mScarlet3). **d,e.** The same experiment as depicted in a,b but using other spectral settings for the red channel: excitation at 562-587 nm and emission at 593-616 nm. **f.** The data of figure e after correcting the red intensities for spectral throughput of the microscope, relative to mScarlet3. The slopes of the linear regression line that is forced the origin was used for calculating the relative brightness. The slope was calculated as ∑RC/∑C_2_ in which R is the RFP intensity value of a single cell (vertical axes) and C is the CFP intensity of the same cell (horizontal axes). The standard error of the slope (indicated in Fig. 3a) was calculated as Sqrt((∑R_2_-(∑RC)_2_/∑C_2_)/((n-1)∑C_2_)). In which n is the number of cells.

**Figure 3.**
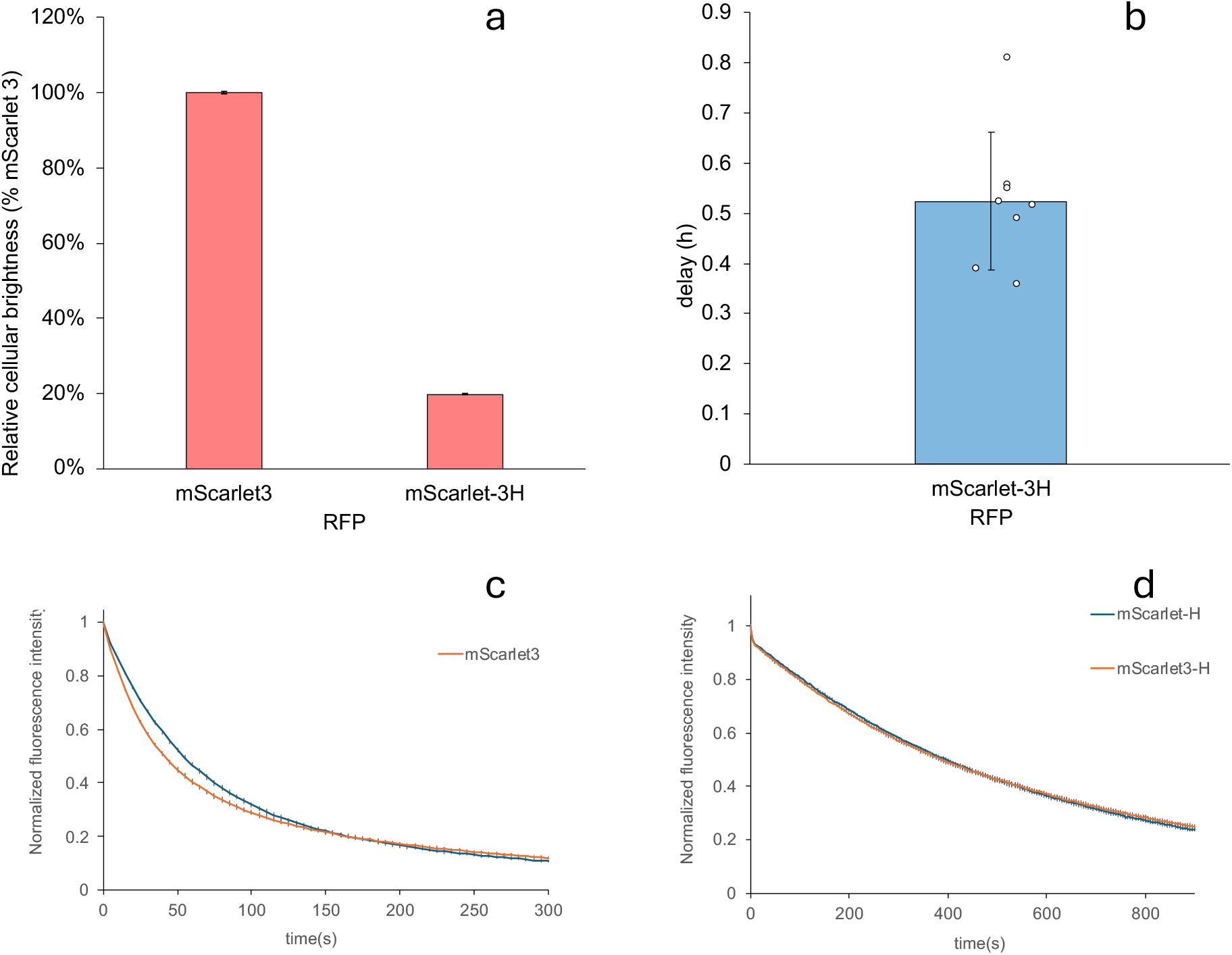
Quantification of cellular brightness, maturation speed and photobleaching kinetics. **a.** Relative brightness (average ± sd) of mScarlet3 and mScarlet3-H in HeLa cells 24 h after transfection, determined by normalized, spectrally corrected ratios of red to cyanfluorescence intensity found in cells upon co-production of RFP and mTurquoise2. **b.** maturation speed of mScarlet3-H in HeLa cells measured as delay (average ± sd in min) relative to co-produced mTurquoise2, n=8 cells. **c.** Photobleaching kinetics of mScarlet and mScarlet3 in living HeLa cells. **d.** Photobleaching kinetics of mScarlet-H and mScarlet3-H in living HeLa cells. For c. and d. cells were subjected to widefield microscopy and illuminated at ∼ 4W/cm2 with 550/15 nm LED power. Only backgroundfluorescence was subtracted and the netfluorescence intensity was normalized to the initial value at t=0. The average ± sd of four individual cells is shown for each curve.

These cellular brightness measurements indicate that mScarlet3-H is comparable in brightness to mCherry (approximately 5-fold less bright than mScarlet3 and mScarlet-I3) (Gadella et al., 2023). By comparing the relative cellular brightness with the relative intrinsic brightness, an estimate can be obtained for the relative extent of maturation. For mScarlet3-H this value is roughly equal to mScarlet3 (109%) 24h after transfection in HeLa cells.

Using the same polycistronic vector and transfection in HeLa cells, a time lapse experiment can be performed in which the accumulated brightness in the cyan and red channel are recorded as a function of time (Gadella et al., 2023). For individual cells, the time traces can be analyzed for a delay between the red and the cyan curves. The maturation delay can be quantified by extrapolating the tangent line fitted at the highest accumulation rates in the two channels towards the horizontal time axis. This yields 2 intercepts in time and providing directly the estimated delay time, a measurement which is independent of detection efficiency or brightness. For mScarlet-3H this yields a maturation delay of 31 minutes (figure 3b). This is comparable to the value published for mScarlet3 (Gadella et al., 2023). So these experiments show that like mScarlet3, mScarlet3-H is a fast and efficiently maturing RFP, similar to mScarlet3. Still the best maturing RFP described to date is mScarlet-I3 with no noticeable delay with respect to mTurquoise2 and a 10% higher overall extent of maturation 24h after transfection (Gadella et al., 2023). It can be concluded that despite efficient maturation in cells, mScarlet3-H displays a greatly diminished overall cellular brightness because of its low intrinsic brightness.

### Photostability

Xiong et al. strongly stressed the marked photostability of mScarlet3-H and suggest it is by far the most photostable RFP described to date (Xiong et al., 2025). In developing several mutant RFPs from mScarlet3, we did not notice that mScarlet3-H had such outstanding performance as compared to mScarlet-H. In fact, we did find other variants with much better photostability (Gadella & van Weeren, unpublised observations). Hence, we carefully revisited photobleaching kinetics experiments and we compared both mScarlet to mScarlet3 and mScarlet-H and mScarlet-3H (see Figure 3c and 3d). In the analysis we only performed intensity-normalization to the initial value, and we did not change the time-axis for photon-flux normalization. Under these conditions, we reconfirm that mScarlet3 bleaches slightly faster than mScarlet (Gadella et al., 2023). Yet we do not see any improvement of the photostabilty of mScarlet3-H over mScarlet-H (see Figure 3d). The curves are perfectly superimposable. Repetitive measurements show the same. So we cannot confirm the claims made by Xiong et al. that mScarlet3-H is significantly more photostable than mScarlet-H (Xiong et al., 2025), nor the qualification of ‘*exceptional photostability*’ (Campbell, 2025) of mScarlet3-H: there is no improvement whatsoever. A difference between our study and the study by Xiong et al. is that they do brightness corrections to the time-axis, whereas we do not correct the time-axis. Furthermore, Xiong et al. only described bleaching of purified proteins or of fixed cells, whereas we report on bleaching kinetics in live cells at 37 degrees. Of course, the latter situation is the more relevant environment for live cell imaging applications. Another issue for mScarlet-H and mScarlet3-H is that they show signs of photochromism: an almost instantaneous loss of 10% of the initial intensity followed by a much slower decline. This resembles photochromism of TagRFP and mApple (Gadella et al., 2023) and likely reflects dynamic photocycling between light and dark states that apparently protect the chromophore against irreversible photobleaching. mScarlet and mScarlet3 on the other hand display ‘normal’ continuous photobleaching. The bleaching kinetics becomes slower when mScarlet(3) is bleached to less than 20%. We may note that (under similar imaging conditions) even at that high bleaching percentage, mScarlet3 still emits more photons than mScarlet3-H in view of its ∼5-times higher brightness. Another important notion is that the excitation power by which we bleach is not at all relevant for typical live cell imaging. For the latter, exposure power is usually 10-50 times lower, and secondly, we only record the emitted light 0.2% of the time that the excitation light is on during photobleaching. Even within these 10 ms integration times we almost saturate the CMOS-based camera detector with normal RFP producing cells. Therefore, one can easily record timelapse movies with 1000 frames and high signal to noise with less than 10% bleaching using cells expressing mScarlet3 and the same widefield microscopy setup. For confocal timelapse imaging, we already described 24 h continuous imaging of actin filaments in live cells without noticeable photobleaching at low excitation power and sensitive photon counting (Gadella et al., 2023). Hence, for regular timelapse live cell imaging, a photostability higher than we see for mScarlet3 is actually not required.

### Nanosecond fluorescence decay analysis

Solutions of purified mScarlet3 and mScarlet3-H were analyzed with time-correlated single photon counting microscopy to analyze the fluorescence decay kinetics. As can be seen in figure 4a, mScarlet3 decays with single exponential kinetics and afluorescence lifetime of ∼3.8 ns. mScarlet3-H decays much faster and displays more complex decay characteristics (see Figure 4b). At least 3 exponents are required to accurately describe the decay curve. An average lifetime value of ∼1.0 ns was measured, containing lifetime components of 0.56 ns (24%), 1.2 ns (70%) and 3.2 ns (6%). The low lifetime is the result of additional competing dark (non-radiative) decay pathways from the excited state causing both faster decay and a lowered fluorescence quantum yield. The complex triple exponential decay indicates that mScarlet3-H coexists in multiple conformations with distinct decay characteristics. These conformations likely are dynamically interconvertible, but at a timescale slower than the ns fluorescence decay dynamics. The value of 3.8 ns for mScarlet3 is slightly lower than the value of 3.96 ns that we published in (Gadella et al., 2023). The small difference is explained by a different setup that we used (a Leica Stellaris-Falcon system used here, versus a Picoquant TCSPC setup), which can give rise to polarization-based bias of the lifetimes, because both excitation and detector optics show polarization in the Leica Stellaris-Falcon. We attempted to mitigate this by measuring under magic angle conditions, but this still may leave a small lifetime-bias in view of mixed polarization at the object plane. The slightly faster decay is caused by rotational diffusion of mScarlet after polarized excitation. Since we used the Stellaris-Falcon setup for unmixing and pH sensing applications (see below), we included the values obtained for that system for comparison. The Picoquant system is more reliable in terms of lifetime estimation in view of its complete insensitivity to polarization-based artifacts.

**Figure 4.**
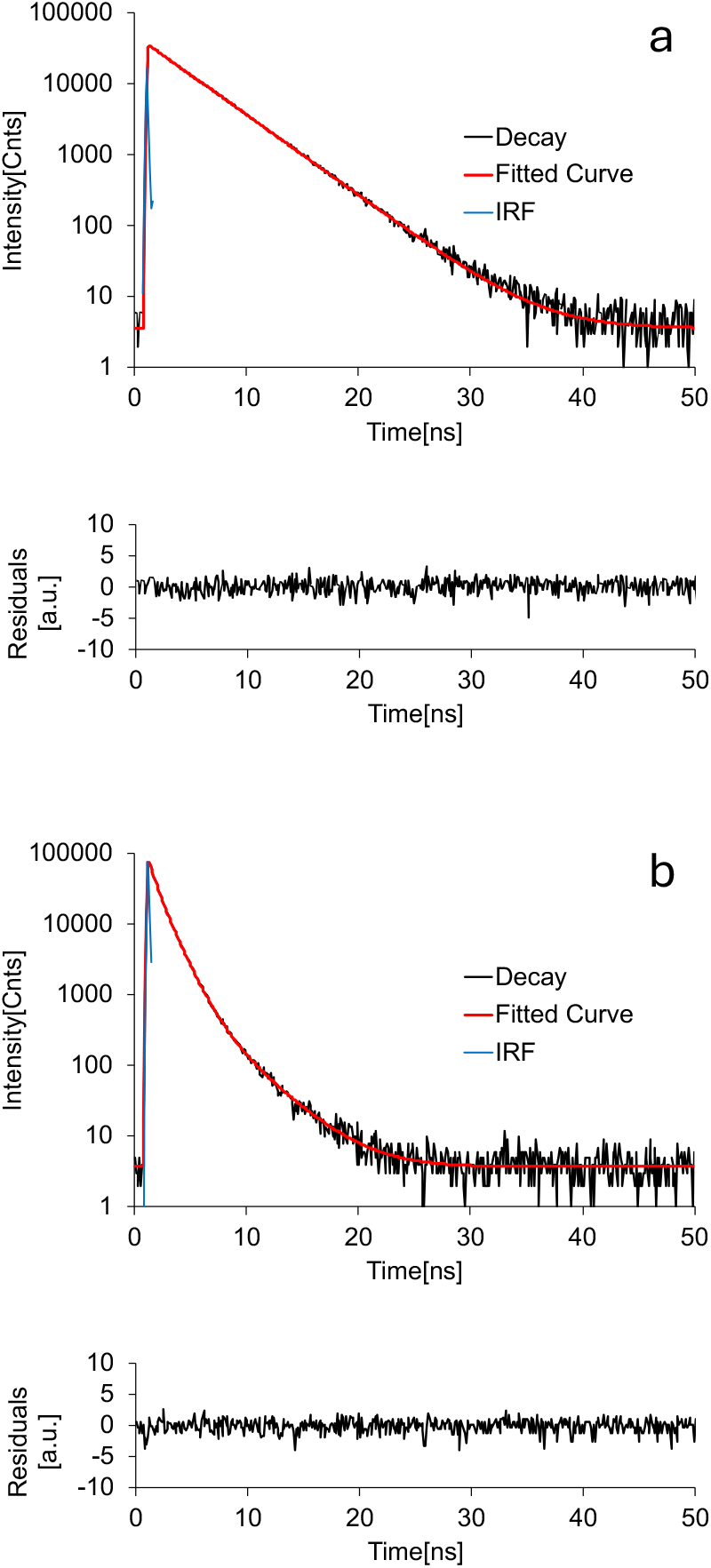
Comparison of time-resolved fluorescence decay of purified mScarlet3 and mScarlet3-H in PBS. **a.** Fluorescence decay of mScarlet3 (top) fitted with a single exponential lifetime of 3.82 ns. The weighted residuals (bottom) show a perfect fit (χ^2^=1.18). **b.** Fluorescence decay of mScarlet3-H (top) fitted with a triple exponential decay with function with lifetimes 0.56 ns (24%), 1.2 ns (70%) and 3.2 ns (6%). The weighted residuals (bottom) show a perfect fit (χ^2^=1.16).

### Fluorescence lifetime unmixing

In view of the much lower averagefluorescence lifetime of mScarlet3-H as compared to mScarlet3, we investigated the possibility of simultaneous imaging of both mScarlet3 and mScarlet3-H-labeled proteins using only a single excitation wavelength and detection channel with FLIM. To this end mScarlet3 was fused to H2B and mScarlet3-H was fused to Lifeact and the constructs were separately expressed in HeLa cells and analyzed by FLIM (see Figure 5a and 5b, respectively). It can be seen that these single-transfected cells display a very homogeneous average lifetime. We see a mean value of 3.5 ns for mScarlet3-H2B and 1.0 ns for mScarlet3-H-lifeact, estimated form the average photon arrival time histograms of the cells (Figure 5a,b). Also a more elaborate TCSPC decay analysis was done of these FLIM data, including a decay and phasor analysis (see Figure 5c-g). Here we see very similar decay characteristics of fusion proteins in living cells at 37 degrees as we see in non-fused purified proteins at room temperature (compare figs 4 and 5). Only the lifetime values are all a bit lower in cells than in PBS solution. This is expected in view of the higher temperature and refractive index in cells as compared to purified protein solutions.

**Figure 5.**
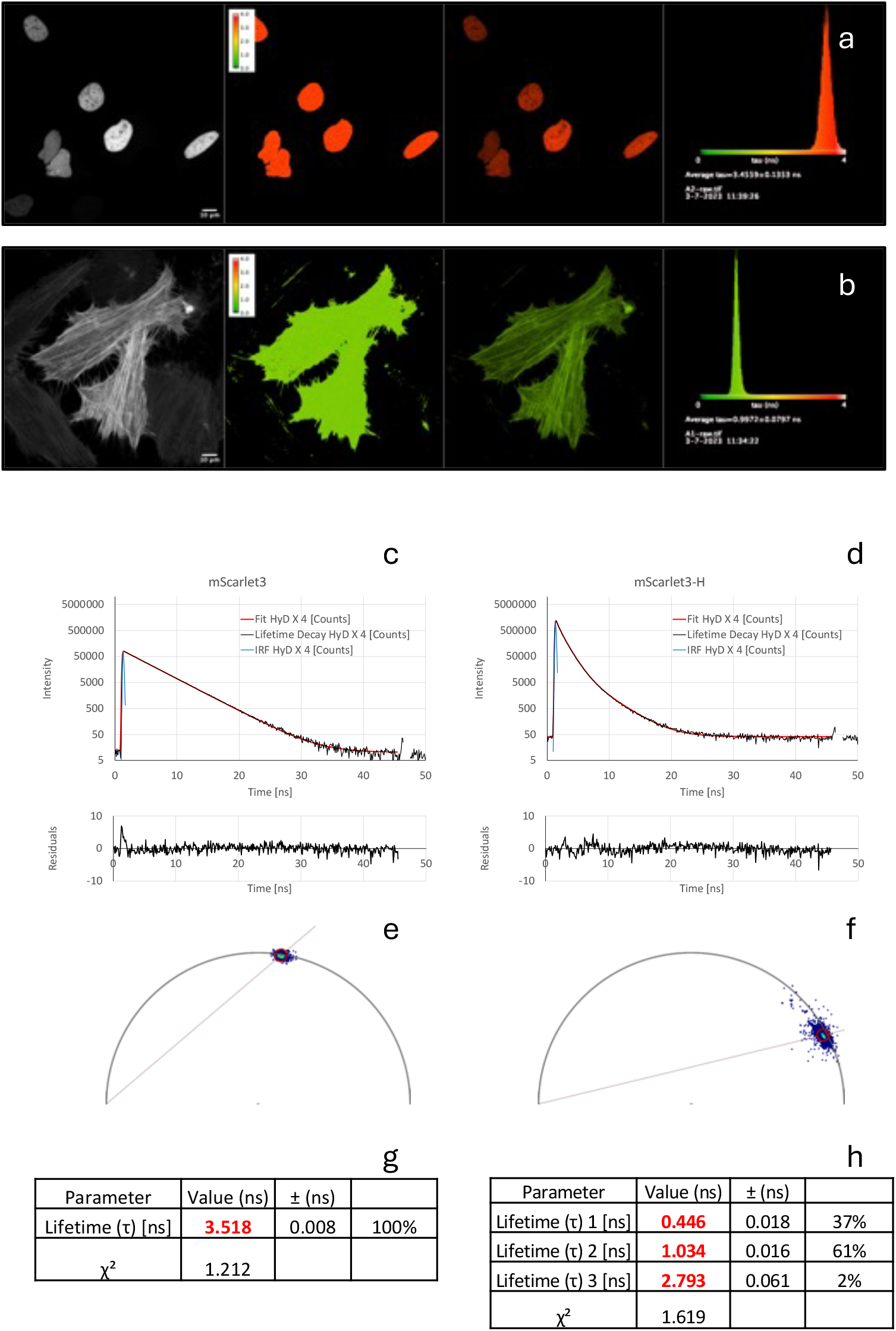
FLIM analysis of fusion proteins of mScarlet3 and mScarlet3-H in live HeLa cells. **a.** Confocal FLIM recording of H2B-mScarlet3 in HeLa cells, from left to right the integratedfluorescence intensity image (bar=10 µm); the average photon arrival time (ns) pseudocolored according to the histogram; the average photon arrival time with brightness according to the measured intensity; the histogram of single pixel photon arrival times scaled between 0 and 4 ns. **b**. Confocal FLIM recording of Lifeact-mScarlet3-H in HeLa cells, with similar layout as 5a. **c.** Fluorescence decay of summed pixels of image 5a (H2B-mScarlet3, top) fitted with a single decay with a lifetime of 3.52 ns. The weighted residuals (bottom) show a perfect fit (χ^2^=1.21). **d.** Fluorescence decay of summed pixels of image 5a (Lifeact-mScarlet3-H, top) fitted with a triple exponential decay function with lifetimes of 0.45 ns (37%), 1.03 ns (61%) and 2.79 ns (2%). The weighted residuals (bottom) show a reasonable fit (χ^2^=1.6). **g,h.** Fit parameters of the decay curves shown in 5c and 5d, respectively

The large lifetime difference between mScarlet3 and mScarlet3-H enables lifetime-unmixing using a single excitation/emission FLIM recording. To illustrate this notion, we cotransfected cells with H2B-mScarlet3 and Lifeact-mScarlet3-H. As can be seen in Figure 6a, we see both the actin cytoskeleton and nuclei in the intensity image within a single FLIM recording (see left panel of Figure 6a). In the average arrival time (or lifetime) image (second panel of Figure 6a) we see a lower value of the photon arrival time in nuclei than in the cytoplasm, indicating lifetime contrast within the FLIM image. In the intensity-weighted colored lifetime image (third panel of Figure 6a) clearly the actin filaments appear as green (low lifetime) whereas nuclei color red (high lifetime). In the corresponding histogram (right panel of Figure 6a) a bimodal distribution is visible with a high abundance of pixels with a ∼ 1 ns average arrival time and fewer pixels with a more widely distributed arrival time around 3 ns. The latter is caused by mixing high and low lifetimes around nuclei where both actin and H2B were visible in the recorded confocal image. In Figure 6b, the phasor diagram is shown of the FLIM recording of Figure 6a, and the coordinates (phasors) for pure mScarlet3-H2B (red circle) and mScarlet3-H-Lifeact (green circle) were copied from Figure 5e and 5f. It can be seen that all pixels in the image have phasor coordinates along a line connecting the two pure phasors. Consequently, the pure phasors can be used to assign single pixel fluorescence intensity values to the two phasor coordinates according to the leverage rule, since the phasor diagram is linear with respect to steady-state fluorescence intensities (Digman et al., 2008). In figure 6c, the resulting unmixed composite color image is shown with mScarlet3 shown in red and mScarlet3-H in green. Also the individual unmixed images are shown in Figure 6d and 6e. These unmixed images clearly demonstrate that the lifetime difference between mScarlet3 and mScarlet3-H can be employed for single-channel lifetime-unmixing, since the nuclear localized mScarlet3 can be completely separated from the actin filaments localized mScarlet3-H (figure 6 d,e). Lifetime unmixing of spectrally similar FPs has been described before (Goedhart et al., 2010; Kremers et al., 2008) but this was not yet described for unmixing lifetime images of live cells labeled by two red fluorescent proteins variants with different lifetimes to our best knowledge. In principle, multiple different detection channels (cyan, green, yellow, red) with the simultaneous use of multiple sets of spectrally similar FPs with different lifetimes would allow further multiplexing options.

**Figure 6.**
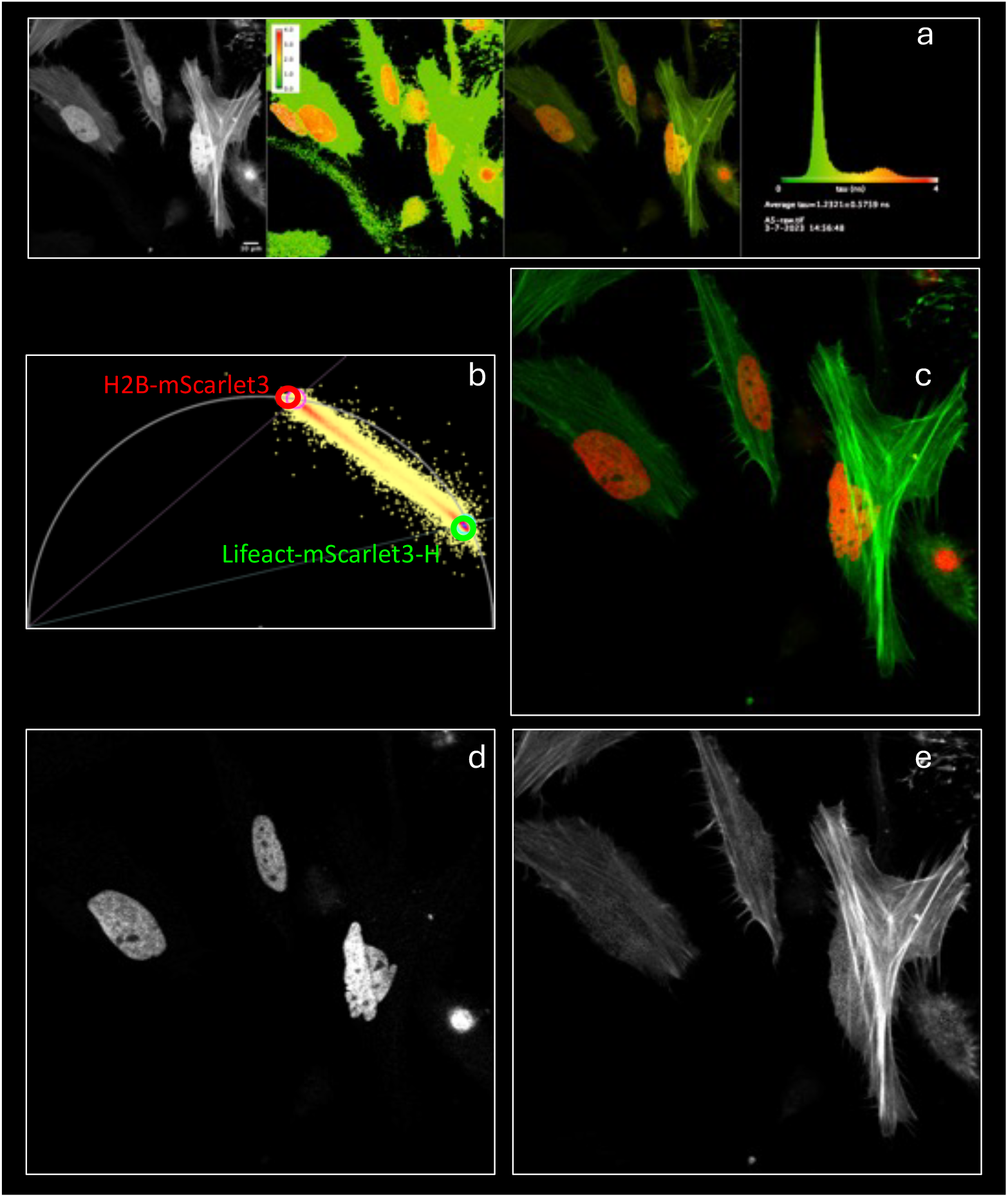
FLIM unmixing of mScarlet3 and mScarlet3-H in live HeLa cells. **a.** Confocal FLIM recording of H2B-mScarlet3 and Lifeact-mScarlet3-H cotransfected in HeLa cells, from left to right the integrated fluorescence intensity image (bar=10 µm); the average photon arrival time (ns) pseudocolored according to the histogram; the average photon arrival time with brightness according to the integrated intensity; the histogram of single pixel photon arrival times scaled between 0 and 4 ns. **b**. Phasor plot of the experiment shown in 6a (analysis at second harmonic, corresponding to 40 MHz) with indicated coordinates corresponding to the phasors of pure H2B-mScarlet3 (red) and Lifeact-mScarlet3-H (green) that were obtained from figure 5 and used for unmixing. **c**. Unmixed composite intensity image with green intensity attributed to the mScarlet3-H phasor and red intensities attributed to the mScarlet3 phasor. **d,e.** Individual unmixed intensity images attributed to the mScarlet3 phasor (**d**) and the mScarlet3-H phasor (**e**).

### pH dependence

As noted by (Xiong et al., 2025), mScarlet-3H has a funny pH-dependency. It becomes brighter at lower pH, like mScarlet-H. Yet at very low pH it becomes dimmer. In figure 7, we further investigated the pH dependence, and we found that besides intensity changes there are also spectral changes: the emission spectrum shifts to higher wavelengths at lower pH and large intensity differences can be seen for mScarlet3-H. For comparison, we also included mScarlet3. Whereas mScarlet3 shows a gradual change in spectra at acidic pH (due to its pK of 4.5, (Gadella et al., 2023)), mScarlet3-H has a peculiar intensity maximum at pH 5, accompanied with a red-shift (compare Figure 6a-f). As stated by Xiong et al, the quantum yield of mScarlet-3H is increased at lower pH and they provide data that suggest simultaneous decrease of the absorbance (Xiong et al., 2025). The increased quantum yield would suggest also an increased fluorescence lifetime. For that reason, we also performed afluorescence lifetime analysis of purified mScarlet3 and mScarlet3-H as a function of pH (see Figure 7). It can be seen that mScarlet3 has high average fluorescence lifetimes over the entire pH range (2-12), mScarlet3-H has a large reduction in fluorescence lifetime at physiological pH and higher (lifetime of ∼ 1 ns), but at acidic pH an increase in fluorescence lifetime can be observed (to ∼3 ns). The pK (or pH for half maximal increase) of mScarlet3-H is ∼ 6 (see Figure 7a). For mScarlet3, the pK for the small lifetime decrease at acidic pH is lower, ∼5. The pK of 6 for mScarlet3-H is quite close to physiological pH. Yet the fluorescence intensity does first increase at (towards pH 5) and then at more acidic pH start to decrease. This indicates that at very acidic pH the decrease in absorbance is much stronger than the quantum yield increase, leading to an overall decreased molecular brightness below pH 5. The increased absorbance and fluorescence lifetime at pH 5 suggests the formation of a conformation of the mScarlet3-H chromophore closer to that of mScarlet3 (which displays a higher extinction coefficient and higher quantum yield). At low pH the chromophore of mScarlet3 and mScarlet3-H become protonated and this leads to a rapid loss of fluorescence intensity (due to absorbance loss). Presumably, the pK for protonation of the chromophore itself is similar for both proteins at pH ∼4. The complex pH characteristics of mScarlet3-H, with two partially antagonistic acting pKs one for lifetime and red shift at pH 6 and one for chromophore protonation at pH ∼4, should be taken into consideration for applications in which mScarlet3-H would be used for lifetime unmixing. The long lifetime component of mScarlet3-H at acidic pH can be mis-assigned to mScarlet3. Still at neutral (physiological) pH (e.g. in cytoplasmic or nucleoplasmic environments), no big deviations of thefluorescence lifetime of mScarlet3-H from its low value around 1 ns are expected.

**Figure 7.**
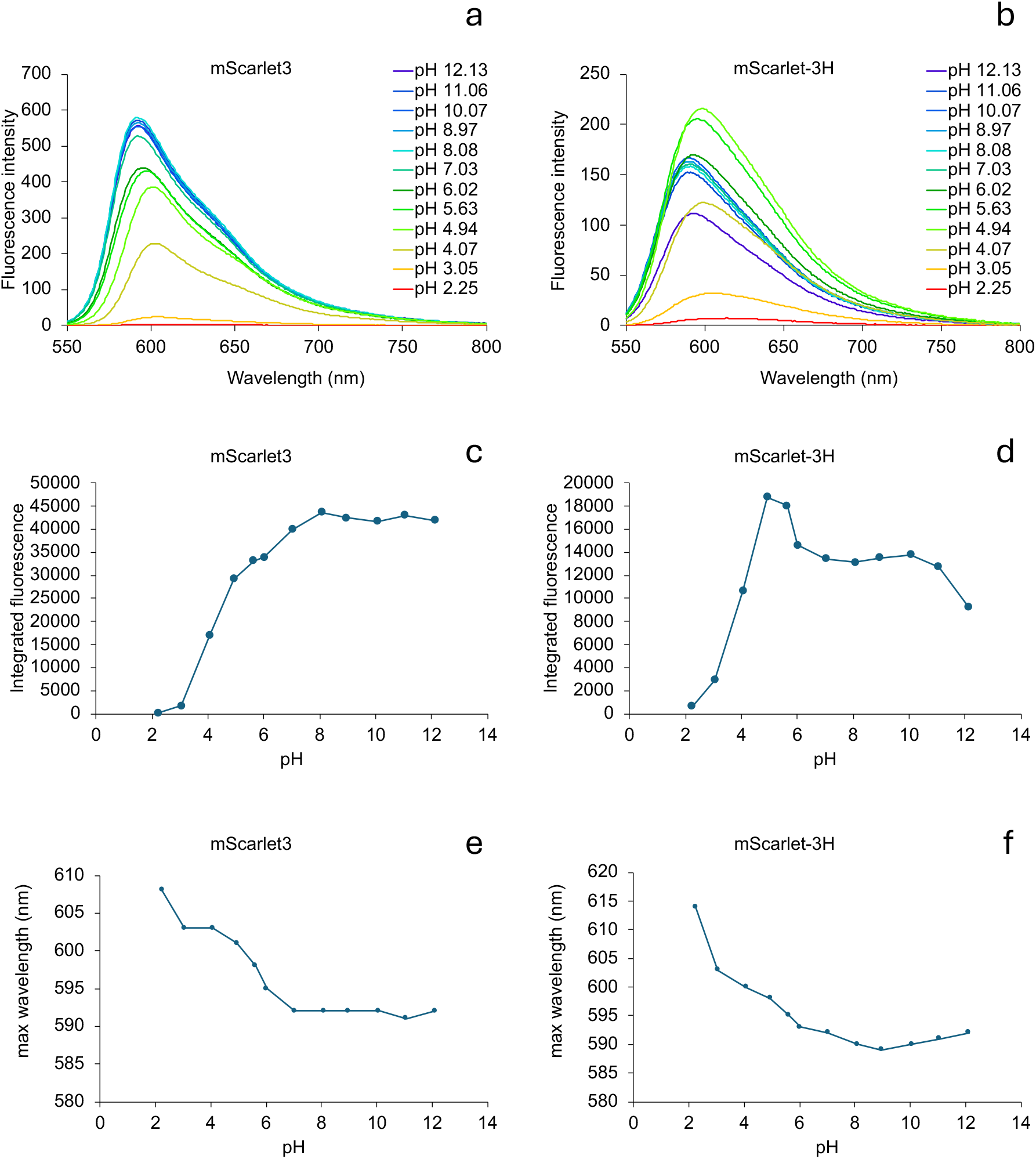
pH dependence offluorescence emission spectra of purified mScarlet3 and mScarlet3-H. **a,b.** Fluorescence emission spectra of mScarlet3 (a) and mScarlet3-H (b) at diberent pH values. **c,d.** Total integratedfluorescence emission of mScarlet3 (c) and mScarlet3-H (d) as a function of pH. **e,f.** Maximalfluorescence emission wavelength of mScarlet3 (e) and mScarlet3-H (f) as a function of pH. Allfluorescence data were taken with 540 nm excitation. Both excitation and emission slits were set at 5 nm and spectra were corrected for backgroundfluorescence (recorded with blanc bubers) and instrument response.

### mScarlet3-H as lifetime based-pH sensor

The increased fluorescence lifetime at acidic pH can also be utilized as an advantage, since it allows to use mScarlet3-H as a quantitative fluorescence lifetime-based pH sensor in this acidic pH range. To that end, as summarized in figure 7b, we applied phasor analysis of the fluorescence decay of both mScarlet3 and mScarlet3-H. For mScarlet3, it is clear that between pH7-12 it behaves as a long lifetime monoexponential decaying species with a phasor coordinate at the semicircle. At pH 6 only a very marginal shift is observed. And at lower pH we see a small right-way phasor trajectory towards lower lifetimes also inwards into the semicircle towards pointing to slightly heterogeneous decay. mScarlet3-H at pH 8-10 occupies a coordinate inwards with respect to the semicircle at a much lower lifetime. Then upon decreasing the pH from 8 to 4 a remarkable linear phasor trajectory is observed corresponding to average lifetimes between 1 and 3 ns. This clearly points to a single pK effect with two species: one with a high lifetime (∼ 3 ns) and one with a low lifetime (∼ 1 ns) at different abundance depending on pH in this region. The halfway transition is at pH 6, which would be a fairly accurate estimate of the pK (also consistent with Figure 7a). The linear trajectory underlines that mScarlet3-H can be used as a genetically encoded quantitative fluorescence lifetime-based pH sensor. The negative lifetime-pH response of mScarlet3-H is quite unique, because most red and green fluorescent proteins are not acid tolerant and become less bright with decreasing pH. The use of fluorescent protein-based pH indicators is already quite established (Miesenböck Gero, 1998). Also red fluorescent protein pH sensors have been described (Li & Tsien, 2012; Shen et al., 2014), including the mScarlet-derived pHmScarlet (Liu et al., 2021). Still most FP-based pH sensors are ratiometric or intensiometric. Recently also fluorescence lifetime-based pH sensing was described based for the red fluorescent protein mApple (Rennick et al., 2022) and for mScarlet (Lazzari-Dean et al., 2022). Most FP-based pH sensors have pH optima near neutral (Li & Tsien, 2012; Liu et al., 2021; Shen et al., 2014). A disadvantage of most (red)FP-based pH-sensors is that they become much less bright at acidic pH, which makes them less sensitive in acidified compartments. This complicates the use RFP-based pH-sensors for the study of pH in acidified endomembranes. mScarlet3-H, however, displays a negative response: its lifetime increases at lower pH and it is less bright at neutral pH. The FLIM calibration shows there is a huge lifetime dependency in a narrow pH range (pH 5-pH 7) a range which is highly relevant for endomembrane compartments.

A notorious misunderstanding in literature is that some RFPs form aggregates in cells, which is concluded from appearance of punctate structures after prolonged expression of (un)fused RFPs in the cytosol. While some RFPs are not monomeric, these punctate structures in most cases are not aggregates within the cytoplasm but lysosomal structures (Katayama et al., 2008; Shemiakina et al., 2012). This process involves autophagy of the cytoplasm and reflects a normal homeostasis in which continuous removal of ‘old’ cytoplasmic material is in balance with production of new protein material through synthesis on ribosomes. Since RFPs display a lower pK and are more resilient to partial proteolysis than most GFPs, this process was not visible with *Aequorea victoria* derived GFPs or YFPs (Katayama et al., 2008). We noticed that cells expressing unfused mScarlet3-H showed accumulation within punctate structures that are clearly visible 72 h after transfection. We analyzed these cells with FLIM and we could see a low lifetime (∼ 1 ns) in the cytoplasm and nucleoplasm (see Figure 9a, corresponding to the regular mScarlet3-H lifetime also seen 24 h after transfection), but and an elevated lifetime in the punctate structures. By superimposing the calibrated phasor trajectory of Figure 8b, we infer that the nucleoplasm and cytoplasm have a neutral pH value, but the punctate structures have a reduced pH between 7 and 6 (Figure 10b). This lifetime change indicates indeed mScarlet3-H accumulates in the lumen of a membrane-bounded endocytic/lysosomal environment and does not reflect aggregation within the pH-neutral cytoplasm. It again underlines the functional contrast mScarlet3-H gives about the local pH.

**Figure 8.**
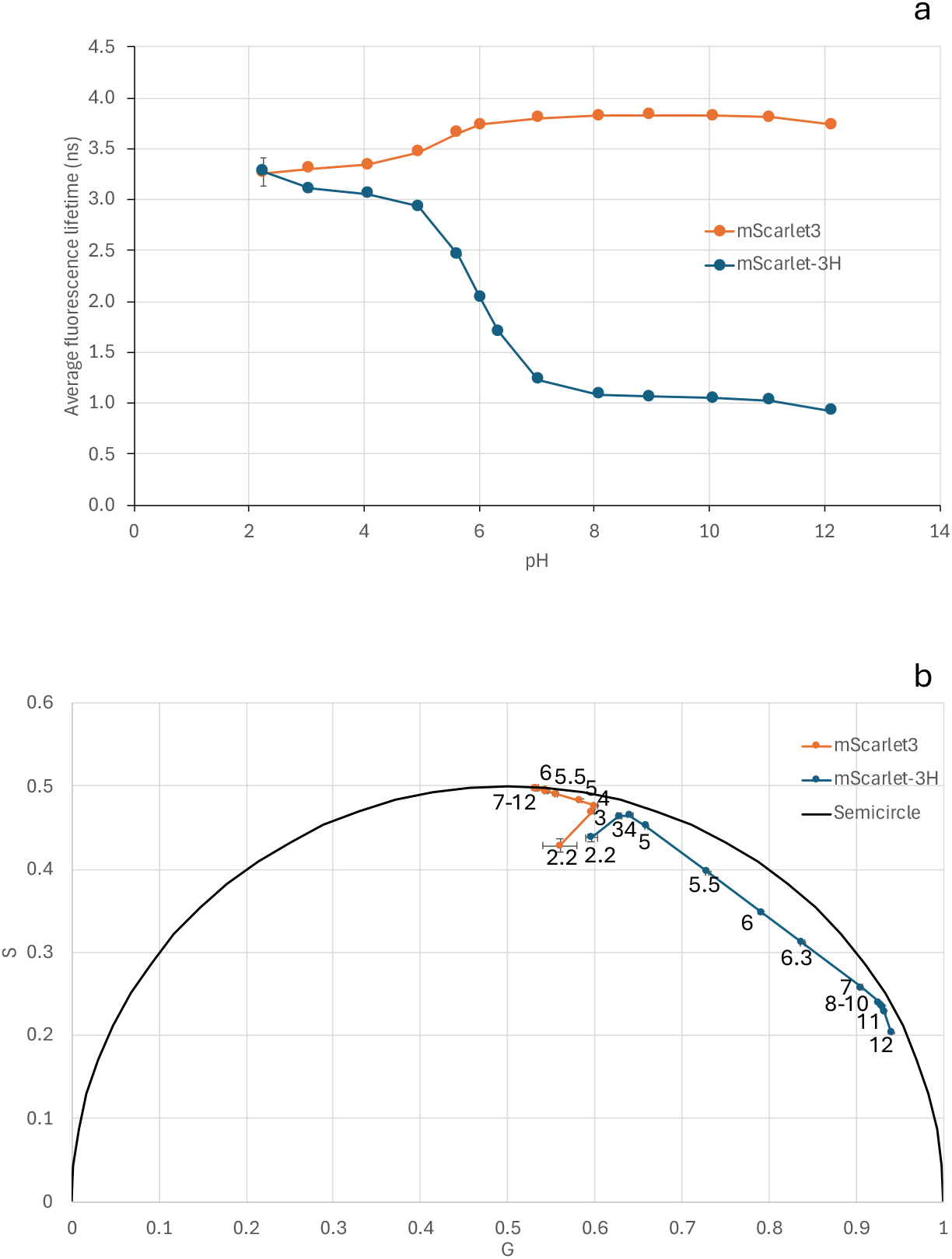
pH dependence of fluorescence lifetimes of purified mScarlet3 and mScarlet3-H. **a.** Intensity-weighted average fluorescence lifetimes of mScarlet3 and mScarlet3-H at diberent pH values. **b.** Phasor representation offluorescence lifetime analysis for mScarlet3 and mScarlet3-H at diberent pH values. Note remarkable linear trajectory for mScarlet3-H between pH 4-8, and pH independence for mScarlet3 between pH 7-12. All data were recorded using 561 nm @ 20 MHz excitation and 590-650 nm emission.

**Figure 9.**
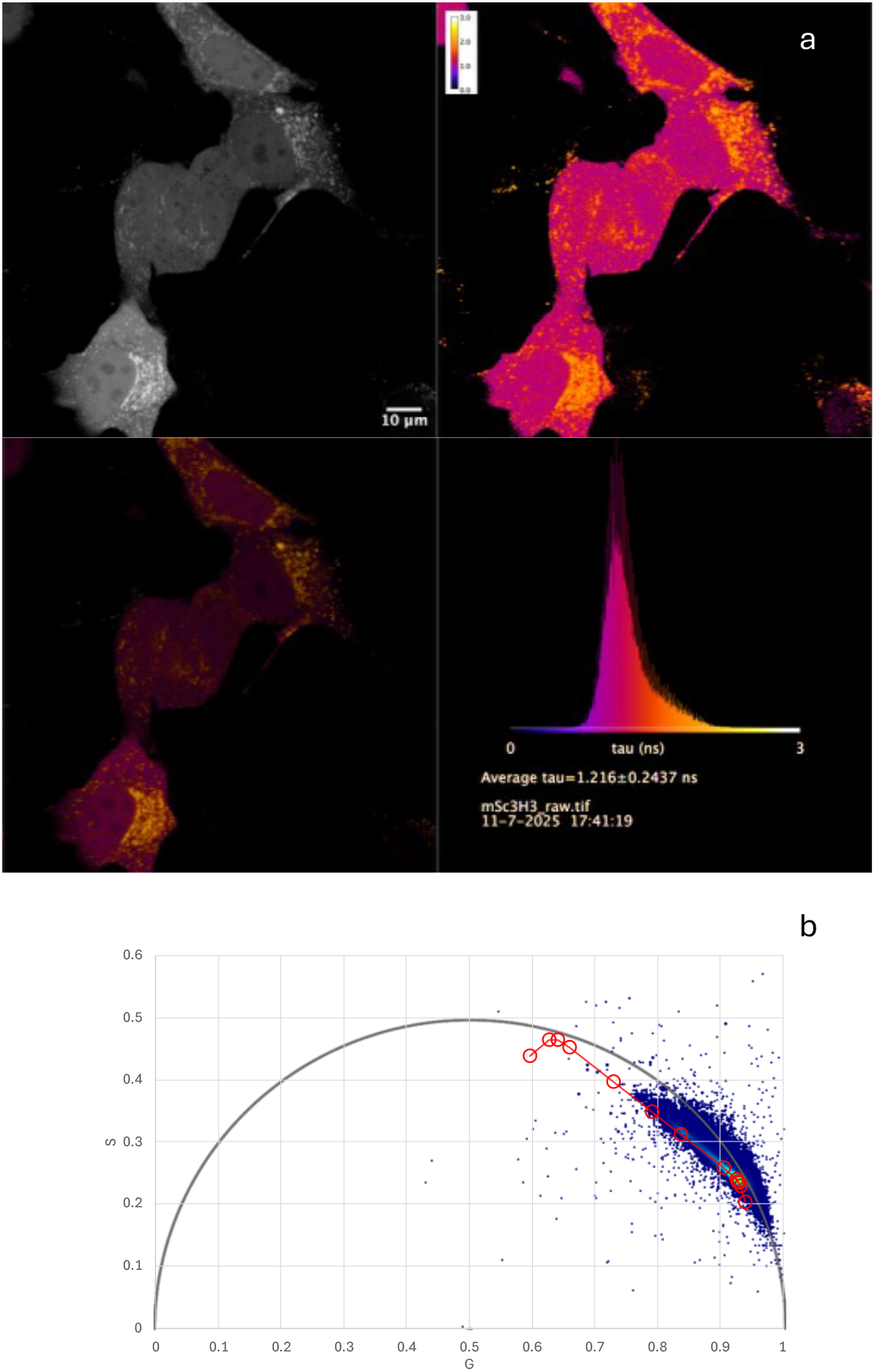
Removal of cytoplasmic mScarlet3-H by autophagy as visualized by its pH-dependent fluorescence lifetime in live cells. **a.** Fluorescence lifetime image of (unfused) mScarlet3-H expressed in live HeLa cell, 3 days after transfection during which ‘old’ cytoplasm is removed by autophagy. From left to right and top to bottom: intensity image (enhanced with gamma=0.5 to better show less intense regions); the average photon arrival time (ns) pseudocolored according to the histogram; the average photon arrival time with brightness according to the measured intensity; the histogram of single pixel photon arrival times scaled between 0 and 3 ns. **b.** Phasor analysis of the FLIM recording of Figure 10a. Superimposed is the phasor calibration for mScarlet3-H in pH buber solutions as shown in Figure 8b. This reveals the neutral pH of the remaining nucleoplasmic and cytoplasmic localized mScarlet3 and accumulation of mScarlet3-H in acidified autophagosomes with a pH between 7 and 6.

## Conclusions

In comparing our characterization of mScarlet3-H with the recently published results by (Xiong et al., 2025), we noticed a few differences. The molar extinction coefficient reported by (Xiong et al., 2025) of 102685 M^-1^cm^-1^ is 23% inflated as compared to our estimate (of 79,040 M^-1^cm^-1^) and inconsistent with other mScarlet variants with blue-shifted spectra carrying the M163H mutation (Bindels et al., 2017). Our estimates of the mScarlet-3H quantum yield at neutral pH and the maturation speed are very similar to the values published by (Xiong et al., 2025). As a consequence, we propose a molecular brightness of 14.1. It is of note that several parameters by Xiong et al. are proportional to the molecular brightness (including the normalized bleach time) and, therefore, may be inflated as well. The diminished extinction coefficient and the much lower quantum yield make mScarlet3-H a less suitable FRET acceptor than mScarlet3, especially in ratiometric FRET applications where the much lower sensitized emission (proportional to the quantum yield) of mScarlet3-H contributes to much reduced ratiometric contrast (Gadella et al., 2023).

For sensitive live cell imaging, the 5-fold brighter mScarlet3 and mScarlet-I3 are markedly preferred over mScarlet3-H. Especially in case of low or endogeneous expression levels, brightness is very important. One might increase the excitation power for mScarlet3-H five times to get an equal intensity as for mScarlet3, but this also will increase the background autofluorescence levels and phototoxicity. Also the very low pH-dependency of mScarlet3 between pH 6-12 make it a reliable fusion tag with producing fluorescence intensity according to its local concentration.

The large lifetime contrast between mScarlet3 and mScarlet3-H enables lifetime-based unmixing in single FLIM recordings. This feature requires a pH between 7-10 to work well. In combination with other lifetime variants of different FP spectroscopic classes, multiplexing applications are possible. Lifetime unmixing allows 2 channel recordings by using only a single excitation laser and single detector channel. Besides options for further multiplexing, this has advantage over regular two-channel recording employing twofluorophores of different spectral classes. For regular 2-channel recording, two different excitation lines are required, with drawbacks related to phototoxicity. In addition, for the blue shifted channel, typically quite narrow bandpasses are required to avoid bleedthrough, which is suboptimal in terms of detection efficiency. For lifetime unmixing, the entire emission spectrum can be used. Another possible application of lifetime-based unmixing is that the unmixing is proportional to the steady state fluorescence intensity contribution of each pure species according to the leverage rule (Digman et al., 2008). So when the relative intrinsic brightness is known for both fluorophores, this intensity ratio can be converted into a quantitative molecular abundance ratio at every pixel of an image. To extract such quantitative information with two lasers and two detection channels withfluorophores from different spectral classes is extremely diNicult and would require extensive instrument (including laser-intensity) calibrations and controls.

mScarlet3-H with its pH sensitivity, enables sensitive phasor-based pH imaging with FLIM in a pH region between 5-7. Previously mApple was described as lifetime-based RFP pH sensor (Rennick et al., 2022). mScarlet3-H is preferred for applications in endomembranes in view of its lower pK around 6 as compared to mApple with a pK of 7. Furthermore, the negative pH response (both lifetime and intensity) of mScarlet3-H yield a higher sensitivity at low pH than for mApple which becomes dark at lower pH. Finally, in view of its extreme photochromicity, great caution should be taken with any quantitative microscopy with mApple in general (Gadella et al., 2023; Shaner et al., 2004). Recently, a preprint was published describing lifetime-based pH sensing with mScarlet which shows a decrease in fluorescence lifetime from pH 6 to 4 from 3.5 ns to 2.2 ns and a pK around 5. They describe a low pH in lysosomes using a LAMP-mScarlet fusion (Lazzari-Dean et al., 2022). We see a similar pK for mScarlet3 (figure 7) but a much smaller lifetime decrease at acidic pH, whereas we see a higher pK for mScarlet3-H, and a much larger pH-dependent lifetime variation. A large advantage of fluorescence-lifetime based sensors is that they are much more robust and quantitative than ratiometric or intensiometric sensors because they do not depend on excitation power, expression level, detection efficiency, emission filtering or light loss due to light scattering or inner filtering (van der Linden et al., 2021). This also allows for deep tissue quantitative pH imaging using mScarlet3-H. Because of the benefits of lifetime-based sensing, several new single fluorescent protein-based genetically encoded sensors have been reported recently. These lifetime-sensors directly confer a conformation change into a fluorescence lifetime change without involving FRET. Currently the palette includes lifetime-based sensors for [Ca2+] (van der Linden et al., 2021, 2025), lactate (Koveal et al., 2022), redox state (Rosen et al., 2025), ATP, cAMP, citrate and for glucose (Zhong et al., 2024). With mScarlet-3H, pH (between 4-8) can be added to that list. Furthermore, since all other lifetime sensors are based on cyan or green emitting FPs, coexpression of these with mScarlet3-H allows simultaneous ion or metabolite and pH imaging with dual lifetime imaging recordings.

Another advantage of mScarlet-3H (as of all mScarlet variants) is that they are cysteine-free proteins and hence do not suffer from cysteine-related redox issues in endomembrane compartments. This would make mScarlet3-H a highly suitable genetically targetable pH sensor or fusion-tag for studying acidified endomembrane compartments. Autophagy of the cytoplasm can be monitored with the pH-based lifetime contrast. The unequivocal determination of the identity of these lower pH structures would deserve further study by cell biologists specialized in these compartments. Here correlative light and electron microscopy (CLEM) is required, since certain endosomal or lysosomal markers fused to FPs end up in multiple endomembrane compartments (Fermie et al., 2018; van der Beek et al., 2022). Here the enhanced chemical stability and resilience towards EM fixation protocols (Xiong et al., 2025) together with the enhanced fluorescent brightness at low pH and functional lifetime contrast reporting on pH, make mScarlet3-H an ideal functional imaging tag for endomembrane studies.

## Acknowledgments

We are grateful to our colleagues Mark Hink and Ronald Breedijk for maintenance of and support with the advanced microscopes and spectroscopy setups at the van Leeuwenhoek Centre for Advanced Microscopy, and to Joachim Goedhart for proofreading of the manuscript.

## Materials and methods

### General Methods Cloning and Transfection

The mScarlet3 and mScarlet-3H constructs were inserted into either the pDRESS plasmid (Addgene #130509) or the pDX vector—a modified TriEX plasmid enabling fluorescent protein expression in both bacterial (under a rhamnose promoter) and mammalian cells (under a CMV promoter) (Bindels et al., 2020). Cloning was performed using the AgeI and BsrGI restriction sites following previously described protocols. HeLa cells (ATCC CCL-2) were used for mammalian imaging. Cells were grown in 24-well plates with glass bottom (MatTek P24G-1.5-13-F) in DMEM (61965059, Thermo Fisher Scientific) containing 10% fetal bovine serum (10270106, Thermo Fisher Scientific) supplemented with 1% of Glutamax (11574466 Thermo Fisher Scientific) under 7% humidified CO2 atmosphere at 37 °C. Transfection was done with polyethylenimine (PEI; 1 mg/mL in ddH_2_O, pH 7.3; Polysciences 23966) using established procedures (Gadella et al., 2023).

### Mutagenesis

The mScarlet3-H variant was generated from mScarlet3 by site-directed mutagenesis to incorporate the M163H mutation using standard techniques, as outlined in (Bindels et al., 2017).

### Protein Purification

Recombinant His-tagged red fluorescent proteins (RFPs) were purified from *E. coli* using protocols reported by (Gadella et al., 2023).

### Spectroscopy Extinction Coefficient

Purified protein samples were diluted in PBS (50 mM Na_2_HPO_4_/NaH_2_PO_4_, 137 mM NaCl, 2.7 mM KCl, pH 7.4) to a final volume of 1 mL. Absorbance spectra were collected with a Libra S70 spectrophotometer (Biochrom) over a 260–700 nm range at 1 nm resolution. A PBS blank was used as reference. To denature the proteins, 100 μL of 10 M NaOH was added, mixed by pipetting, and spectra were recorded until the red chromophore peak disappeared, leaving only the green chromophore peak at 457 nm. The concentration of the denatured chromophore was estimated using an extinction coefficient of 44,000 M^−1^cm^−1^ at 457 nm (Gross et al., 2000; Shagin et al., 2004). From this, the extinction coefficient of the red chromophore was calculated at its absorbance maximum, correcting for the 1.1× dilution.

### Quantum Yield

Proteins diluted in PBS were analyzed for absorbance and fluorescence. Absorbance was measured as described above, with three dilutions giving A_5_4_0_ values between 0.005 and 0.05. Fluorescence emission was recorded using a Jasco FP-8500fluorimeter with a red-extended PMT (R928-23), managed via Jasco Spectral Manager v2.15. Excitation was at 540 nm, and emission spectra were collected from 550–800 nm (1 nm step size, 200 nm·min^−1^ scan rate, 2.5 nm slit widths). Quantum yield calculations were performed using mScarlet3 (QY = 0.751) as a reference.

### Fluorescence Lifetime

Lifetime measurements were conducted on purified RFPs diluted in PBS (same composition as above) or pH buffers (see below) using a Leica Stellaris 8 confocal microscope equipped with an HC PL APO 20x/0.75 objective. Excitation was provided by a white light laser (WLL) set to 85% of its maximum power output. The Acousto-Optical Beam Splitter (AOBS®) operated at 5% transmission, delivering 561 nm pulses at a repetition rate of 20 MHz. For all acquisitions, line averaging was set to 16 and scan speed was 200-400 lines/min. To suppress polarization-dependent artifacts during fluorescence lifetime measurements, a polarization filter was placed at magic angle relative to the main excitation polarization direction (42° at our setup). Emission light in the 590–650 nm range was detected using a HyD X4 detector.

For cellular unmixing experiments, 561 nm pulsed excitation (20 MHz) at 1-3% AOBS transmission and a HC PL APO 40x/0.95 CORR air objective were used. The employed emission bandpass was 575-625 nm with a HyDX4 detector. For cellular pH measurements a 590-650 nm bandpass was used. All cellular microscopy measurments were done without replacing medium directly in the 24-wells plate at 37 degrees and 5% CO2 humidified atmosphere.

### pH Sensitivity

A pH series from 2 to 12 was prepared using a universal buffer containing 50 mM each of citric acid, phosphoric acid, and boric acid with 100 mM NaCl. Each buffer was titrated from a 2× stock with 1 M NaOH (Merck 109137) and diluted accordingly. Final pH values were measured 24 hours later at room temperature. Citric and phosphoric acids (Merck #818707 and #563) and boric acid (Sigma B-0252) were used. Fluorescence emission specrta were recorded with the same instrument and wavelengths as described for the quantum yield determination, except for the exciation and emssion slits that were both 5 nm.

### Microscopy-Based Cellular Brightness Assay

RFP brightness in mammalian cells was measured in HeLa cells in (Bindels et al., 2020). Cells were transfected with 50 ng of pDRESS plasmids encoding expression of a 1:1 ratio of mTurquoise2 and the RFP of interest, along with 150 ng carrier DNA, in 24-well glass-bottom plates. After 24 h, fluorescence images were collected using a Nikon Eclipse Ti-E widefield microscope with LEDs at 440 and 555 nm (SpectraX, Lumencor). Bandpass filters (440/20, 550/15 nm) were used for filterting excitation light. For 440 nm excitation, a triple-band cube (MXU74157, Nikon) was used, for 555 nm excitation a quad band cube (MXU 71640, Nikon) was used. Emission was additionally filtered with a 479/40 (CFP channel) or by a 593/46 nm bandpass (for the RFP channel) (all from Semrock) placed in an optical filter changer (Lambda 10-B, Sutter instrument). For the two detected channels cyan and red, the effective excitation and emission bands were cyan: 430-450 nm excitation, 459-490 nm emission, red: 543-558 nm excitation and 570-616 nm emission. Images were captured using an ORCA-Flash4.0 V2 camera. A 10x CFI Plan Apochromat NA 0.45 objective was used. Image acquisition included a 5×5 tile scan (512×511 pixels per tile, stitched to 2253×2253 pixels). LED intensities were set to 10%, (440 and 555 nm), with integration times of 60 ms. The red-to-cyan fluorescence ratio was analyzed using the ratio_96-wells_macro_v12 macro in (Bindels et al., 2020). Corrections for optical throughput were made using excitation/emission spectra and normalized to mScarlet. Red/Cyan ratios were multiplied by 0.985 for mScarlet3 and 0.692 for mScarlet-3H to yield relative brightness values. Experiments were repeated at least three times in different months and cell lines with consistent outcomes. All cellular microscopy measurments were done without replacing medium directly in the 24-wells plate at 37 degrees and 5% CO2 humidified atmosphere

### Maturation Kinetics

Maturation kinetics were measured using the same experimental setup as for brightness. Starting five hours post-transfection, red and cyan images were taken every 15 minutes for 48 hours. ROIs were manually drawn around 8–10 viable cells, showing steadily increasting fluorescence intensity while lacking detectable fluorescence initially. Only viable cells with continuous and interpretable time traces were included. Cells with discontinuous time traces (due to cell division or apoptosis) were discarded. Intensity values were background-corrected, normalized, and fitted with linear regressions to determine the maximal slope. The intercept of this line with the time axis was calculated and the delay between the intercept found for the red and cyan curves were calculated for 8-10 individual cell traces and averaged

### Photobleaching Kinetics

Photobleaching kinetics was performed as described in (Gadella et al., 2023). Transfection, mounting conditions and the microscope used were identical to those described under the ‘Microscopy-based Cellular Brightness’ section. For imaga acquiosition only the RFP channel was recorded bu using a a ×20 Plan Fluor NA 0.5 air objective and the 555 nm LED was set at 100% power. Timelapse recordings were done within the central ROI of 512×512 pixels at 2×2 binning while continuously illuminating. Every 5s an image was acquired with an integration time of 10 ms, corresponding to a duty cycle of 0.2 %. Excitation power at the object plane was 3.96 W/cm2. For analysis ROIs were drawn around viable cells and the average intensity was determined for 4 individual cells for each time point in the timelapse. Also an area without cells was selected from the timelapse movie to estimate the average background fluorescence and camera bias. For every individual cell time-trace this background value was subtracted for the corresponding time points. The subsequent net cellular intensity-time traces were normalized to their initial value. The average value ±sd for 4 cells is indicated in Figure 3c and d.

### Subcellular Localization and lifetime unmixing

pLifeact-mScarlet-3H_N1 was obtained by digestion of pLifeact-mScarlet3_N1 (189767, Addgene) with AgeI and BsrGI to exchange mScarlet3 for mScarlet-3H. HeLa cells expressing H2B-mScarlet3 (pCS2FA-H2B-mScarlet3 (Addgene 224436)) and Lifeact-mScarlet3 were obtained by transfecting cells that were seeded in uncoated glass-bottomed 24 well plates (MatTek P24G-1.5-13-F) with 50 ng plasmid, 150 ng carrier DNA and 2 µg PEI. 24h after transfection, the cells were analyzed with confocal FLIM in culture medium at 37 °C and 5% CO2 atmosphere using the Leica Stellaris-Falcon setup described above.

## Author contributions

LvW performed the mutagenesis, cloning of constructs, transfection, recombinant protein production and cell culturing. TWJG performed the spectroscopic and microscopy experiments, the data analysis and wrote the manuscript.

